# Melanoma cells adopt features of both mesenchymal and amoeboid migration within confining channels

**DOI:** 10.1101/2021.02.12.431031

**Authors:** Sairisheel R. Gabbireddy, Aram J. Chung, Jeremy S. Logue

## Abstract

For metastasis to occur, cancer cells must traverse a range of tissue environments. In part, this is accomplished by cells adjusting their migration mode to one that is best suited to the environment. Melanoma cells have been shown to be particularly plastic, frequently using both mesenchymal and amoeboid (bleb-based) modes of migration. It has been demonstrated that 2D confinement will promote the transition from mesenchymal to bleb-based migration. However, if melanoma cells similarly transition to bleb-based migration in response to 3D confinement, such as within narrow channels, is unknown. Here, using micro-fabricated channels, we demonstrate that melanoma cells adopt features of both mesenchymal and bleb-based migration. In narrow (8 μm; height and width) channels coated with fibronectin, ~50% of melanoma cells were found to use either mesenchymal or bleb-based migration modes. In contrast, the inhibition of Src family kinases or coating channels with BSA, completely eliminated any features of mesenchymal migration. Detailed comparisons of migration parameters revealed that blebbing cells, particularly in the absence of adhesions, were faster than mesenchymal cells. In contrast to what has been previously shown under conditions of 2D confinement, pharmacologically inhibiting Arp2/3 promoted a fast filopodial-based mode of migration. Accordingly, we report that melanoma cells adopt a unique range of phenotypes under conditions of 3D confinement.

## Introduction

Cell migration constitutes a fundamental process during embryonic development, immune surveillance, wound healing, and cancer metastasis. Melanoma cells, which are known to be aggressive, have been shown to utilize a variety of mechanisms for cell migration, including aspects of mesenchymal and amoeboid migration^1^. This is significant as the ability to switch migration modes is likely required by melanoma cells to traverse the diverse physicochemical environments of tissues.

To metastasize, melanoma cells residing in the skin must travel great distances to reach vital organs. In doing so, cells encounter a range of tissue environments, which may have low-to-high levels of adhesion proteins and confinement^2^. Recently, low adhesion and high confinement was shown to induce a phenotypic transition from mesenchymal to fast amoeboid migration. Embryonic, immune, and cancer cells, including from melanoma, were among the cell types tested and observed to undergo this switch^3^. During fast amoeboid migration cells protrude a large, intracellular pressure-driven, stable bleb. Together with friction, a rapid flow of cortical actomyosin in these blebs provides the motive force for cell movement^4^. Because these blebs lead the cell forward, we previously termed this mode Leader Bleb-Based Migration (LBBM)^5^. Moreover, in melanoma cells, we have previously shown that LBBM requires the actin capping and bundling protein, Eps8, and the down-regulation of Src tyrosine kinase activity^5,6^. Thus, the abatement or prevention of melanoma metastasis is likely to require the simultaneous inhibition of multiple migration modes.

Previously, the phenotypic transition to LBBM has been demonstrated under conditions of low adhesion and high confinement^3^. More specifically, using intravital imagining, cells squeezed between other cells in tumors or developing embryos have been observed using LBBM^7,8^. Similarly, using micro-fabricated devices, compressing cells down to a few microns will trigger the switch to LBBM^3,9^. Using these so called 2D confinement approaches has been useful for the study of LBBM, however, whether melanoma cells undergo a similar transition under conditions of 3D confinement is unclear. Moreover, such conditions, may even trigger novel modes of migration. For instance, breast cancer cells confined within micro-sized channels, may use directed water permeation for migration^10^. While, other cells have been shown to require a heightened level of myosin contractility for migration when confined in microchannels^11^. Therefore, fully understanding cell motile behavior requires comparing 2D and 3D methods of confinement.

Here, we identify that melanoma cells adopt features of both mesenchymal and amoeboid migration within micron-sized channels. However, by inhibiting focal adhesion formation, we find that blebbing cells migrate with increased instantaneous speeds and directionality. Whereas, upon inhibiting Arp2/3, melanoma cells may adopt a fast filopodial-based mode of migration. Thus, revealing that 3D confinement elicits a set of cell motile behaviors distinct from 2D confinement.

## Results

### Melanoma cells adopt features of both mesenchymal and amoeboid migration

In order to evaluate the motile behavior of melanoma A375 cells under conditions of 3D confinement, we micro-fabricated channels using standard photo− and soft-lithography procedures^12,13^. As the deformation of the nucleus is rate limiting when cells are confined below 7 μm, our microchannels were kept at a height and width of 8 μm (Fig. 1A-C)^14,15^. Moreover, while many studies have used PDMS channels attached to cover glass, here we have micro-fabricated channels in which all four walls are made of PDMS (Fig. 1B). Thus, cells sense equivalent levels of stiffness and channel coatings on all sides. To simulate channel-like environments in *in vivo*, we first coated channels with fibronectin (10 μg/mL). Using this approach, we found that cells predominantly adopt two phenotypes with equal frequency (Fig. 2A-C). For phenotype 1, cells protrude in the direction of migration, as well as, appear to attach to the substratum by way of focal adhesions (Fig. 2A; *green arrow* & Movie S1). Simultaneously, at the trailing edge of these cells, are numerous dynamic blebs (Fig. 2A-B). Using EGFP tagged Regulatory Light Chain (EGFP-RLC), myosin is found predominantly at each end and not at the sides of cells in microchannels (Fig. 2B). In contrast, phenotype 2 cells display dynamic blebs at both the leading and trailing edges of cells (Fig. 2A-B & Movie S2). Surprisingly, blebbing cells appear to also form focal adhesions, however, these adhesions are not enriched at the leading edge like phenotype 1 (Fig. 2A; *green arrows*). Moreover, some of these focal adhesions appear on the sides/base of blebs (Fig. 2A; *red arrows*). Myosin is found at the base of blebs and is enriched at the leading and trailing edges of cells, similar to phenotype 1 (Fig. 2B). Although mostly distributed at the cell poles, some myosin can also be found along actin fibers spanning the length of the cell (Fig. 2B). Therefore, under these conditions, melanoma cells adopt features of both mesenchymal and amoeboid migration.

**Figure 1.**
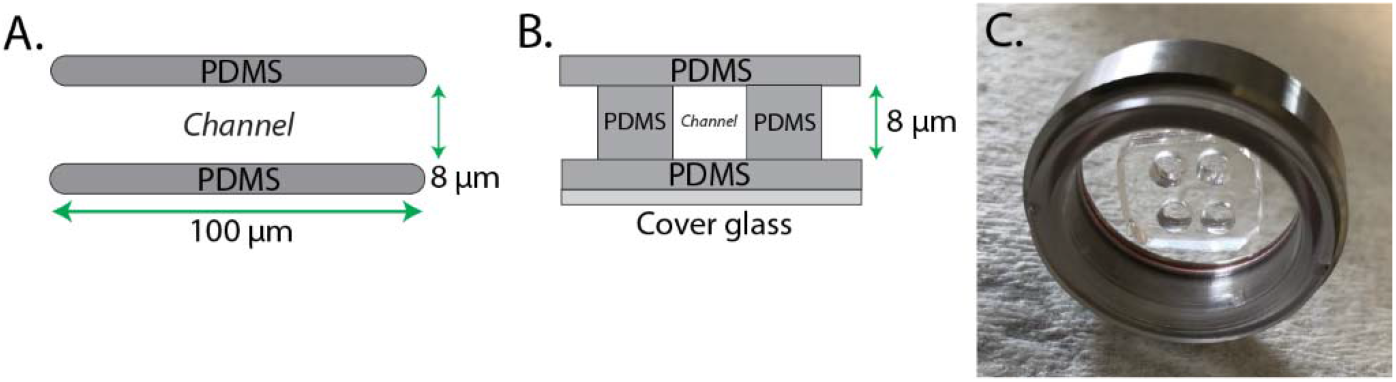
Microchannel design. Throughout this study, cells were confined within 100 μm long (8 μm; height and width) PDMS channels. Microchannels were fabricated with PDMS on all four sides. Top (A) and edge-on (B) views are shown. **C.** Fully assembled microchannel apparatus with two sets of cell loading ports.

**Figure 2.**
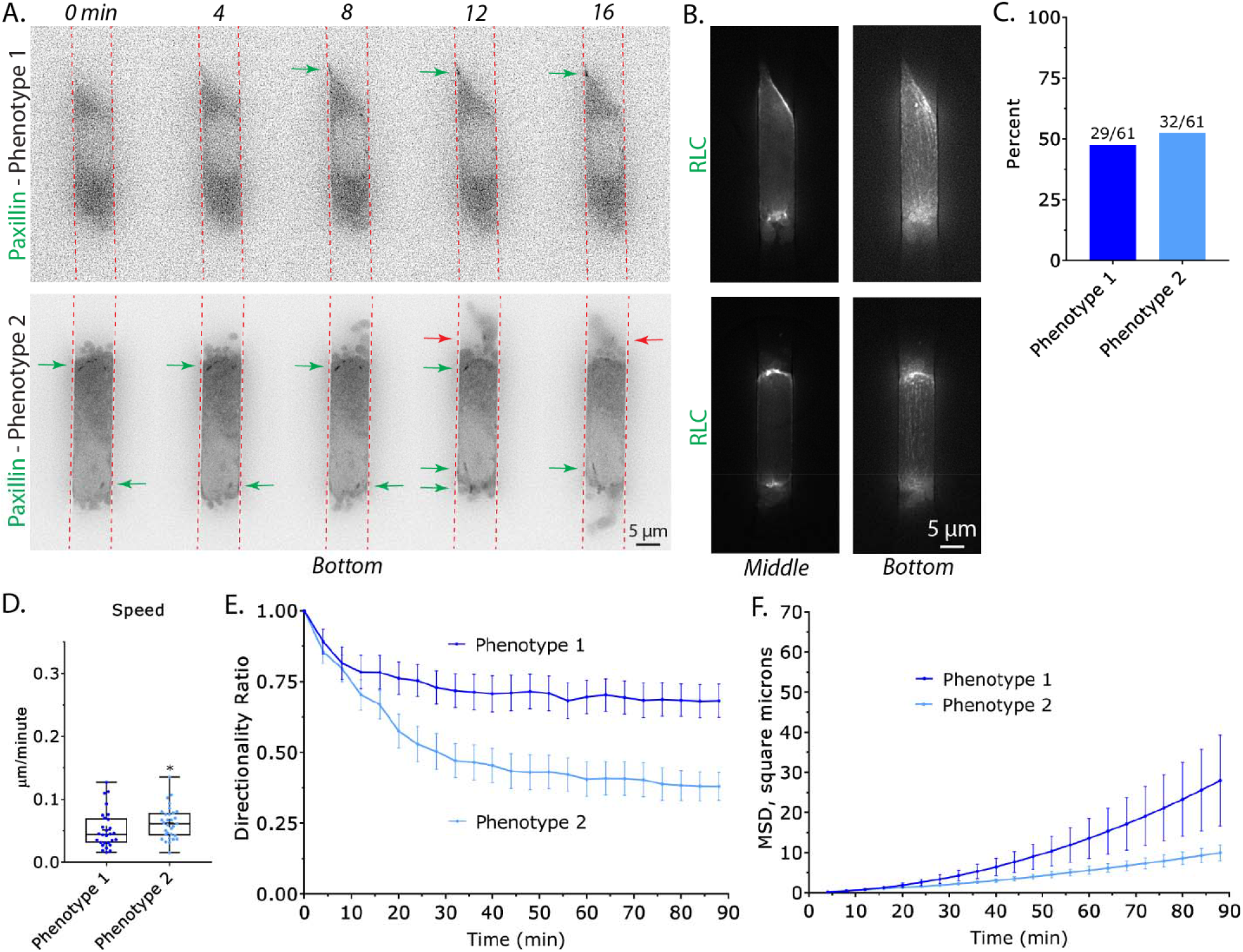
Melanoma cells confined within fibronectin coated microchannels predominantly assume two phenotypes. **A.** Melanoma A375-M2 cells with the focal adhesion marker, EGFP-paxillin, within fibronectin coated (10 μg/mL) microchannels. *Top*, phenotype 1 cells display focal adhesions at their leading edge. Green arrow points to a focal adhesion. *Bottom*, phenotype 2 cells display both focal adhesions and numerous dynamic blebs at their leading and trailing edges. Green arrows point to focal adhesions formed on the cell body. Red arrows point to focal adhesions formed on blebs. **B.** A375-M2 cells with the myosin marker, EGFP-Regulatory Light Chain (RLC), within microchannels for phenotype 1 (*top*) and 2 (*bottom*). Middle (*left*) and bottom (*right*) focal planes are shown. **C.** Percent of A375-M2 cells displaying phenotype 1 or 2 within fibronectin coated (10 μg/mL) microchannels. Cell numbers are shown above each bar. **D.** Instantaneous speeds for phenotype 1 & 2. Statistical significance was determined by an unpaired two-tailed Student’s t-test. **E.** Directionality ratio over time for phenotype 1 & 2. Error is SEM. **F.** Mean Square Displacement (MSD; square microns) over time for phenotype 1 & 2. Error is SEM. All data are representative of at least three independent experiments. * - p ≤ 0.05, ** - p ≤ 0.01, *** - p ≤ 0.001, and **** - p ≤ 0.0001

We wondered how well each phenotype migrated within fibronectin coated (10 μg/mL) microchannels. Accordingly, we manually tracked many cells for 5 hr by time-lapse imaging. Comparing instantaneous speeds, phenotype 2 migrated over 20% faster than phenotype 1 (Fig. 2D). However, the directionality over time of phenotype 1 was found to be significantly greater than phenotype 2 (Fig. 2E). In order to compare the amount of space traveled by each phenotype, we then compared the Mean Square Displacement (MSD) over time for each phenotype. Using this method, cells displaying phenotype 1 were found to travel significantly farther than phenotype 2 over the same period of time (Fig. 2F). Thus, the increased directionality of phenotype 1 cells was found to lead to a significantly higher MSD over time.

### De-adhesion increases the speed and directionality of blebbing cells

Given that melanoma cells adhering to fibronectin continue to bleb, we wondered what effect removing focal adhesions may have on migration parameters in microchannels. Previously, we have used the Src family kinase inhibitor, Dasatinib, to induce the rapid de-adhesion of melanoma cells^6^. In addition, previous studies found that Dasatinib treatment did not impair the ability of these cells to form blebs^6^. Therefore, we used Dasatinib to induce the de-adhesion of cells in fibronectin coated (10 μg/ml) microchannels. Dasatinib treated cells were found to from numerous, prominent, blebs at their leading edge (Fig. 3A; *zoom* & Movie S3). Strikingly, the instantaneous speed of these cells was increased by 2-fold over phenotype 1 & 2 (Fig. 3B). While, the directionality over time of Dasatinib treated cells was between that of phenotype 1 & 2 (Fig. 3C). Likely as a result of their higher instantaneous speeds, Dasatinib treated cells displayed a much larger MSD over time (relative to phenotype 1 & 2; Fig. 3D). In fact, the fastest cells left the channels before the end of the time-lapse (Fig. 3D). Thus, while focal adhesions may increase directionality, the pharmacological inhibition of focal adhesion formation appears to substantially increase the instantaneous speeds of blebbing cells in microchannels.

**Figure 3.**
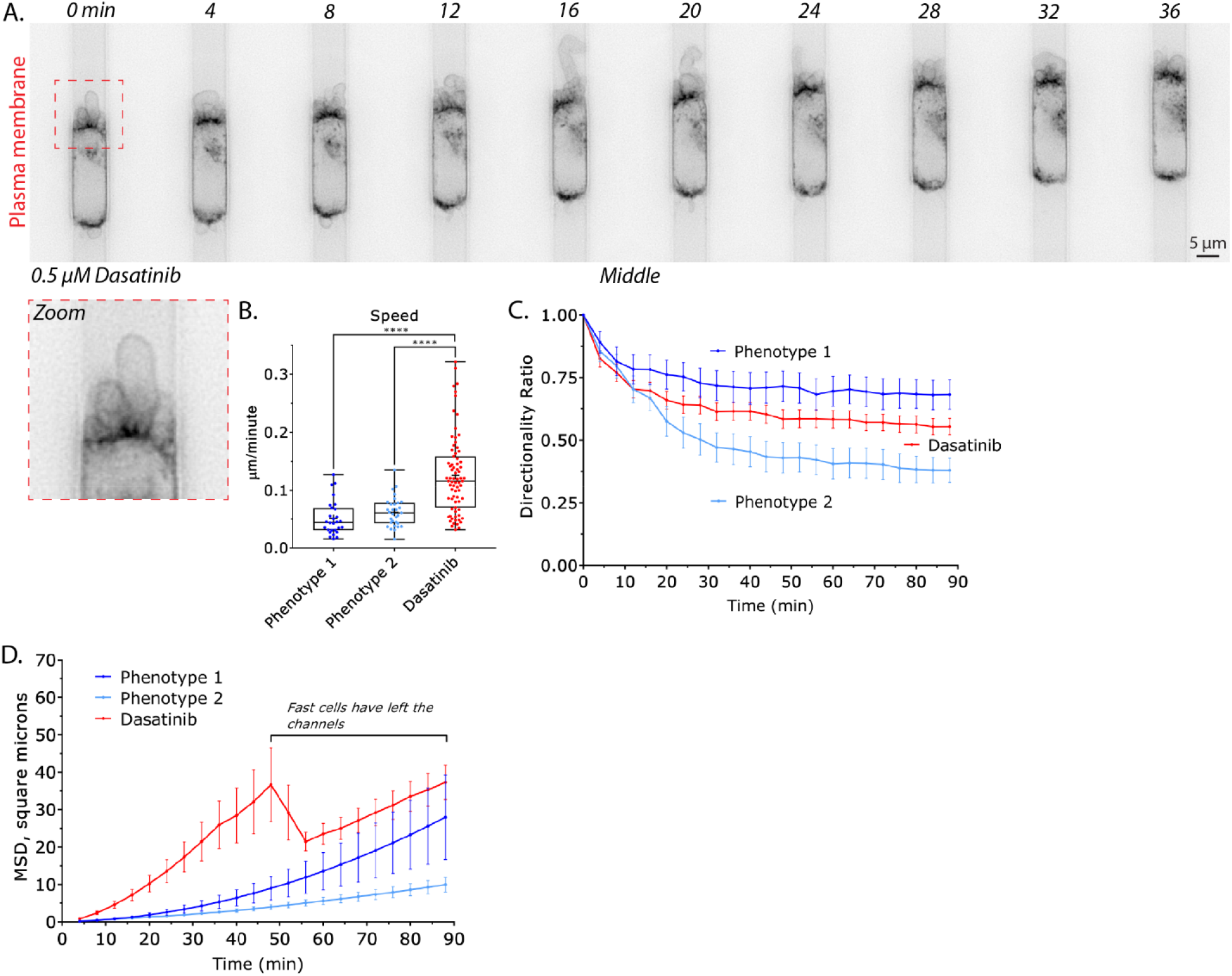
Pharmacologically inhibiting focal adhesion formation increases the speed and directionality of blebbing cells. **A.** Montage of a melanoma A375-M2 cell treated with Src family kinase inhibitor, Dasatinib (0.5 μM), within a fibronectin coated (10 μg/mL) microchannel. Cells were visualized using a far red membrane dye. Zoom shows prominent blebs at the cell leading edge. **B.** A comparison of instantaneous speeds for phenotype 1, 2, and Dasatinib treated cells. Statistical significance was determined by a multiple-comparison test post-hoc. **C.** A comparison of directionality ratio over time for phenotype 1, 2, and Dasatinib treated cells. Error is SEM. **D.** A comparison of Mean Square Displacement (MSD; square microns) over time for phenotype 1, 2, and Dasatinib treated cells. As noted on the right of the graph, a sharp decrease in the graph is the result of fast cells leaving the channels. All data are representative of at least three independent experiments. * - p ≤ 0.05, ** - p ≤ 0.01, *** - p ≤ 0.001, and **** - p ≤ 0.0001

### Low adhesion increases the speed and directionality of blebbing cells

To further clarify the role of focal adhesions in blebbing cells, we passivated microchannels with 1% BSA (Fig. 4A). As BSA is not ligated by integrins, melanoma cells in these channels do not form focal adhesions (Fig. 4B). Similar to the effects of Dasatinib, these cells formed numerous, prominent, blebs at their leading edge (Fig. 4C; *zoom* & Movie S4). Comparing instantaneous speeds, cells in BSA coated microchannels were more than 30% faster (relative to phenotype 1 & 2; Fig. 4D). Like Dasatinib treated cells, the directionality over time of these cells was between that of phenotype 1 & 2 (Fig. 4E). Whereas, the MSD over time of cells in BSA coated microchannels was greater than phenotype 1 & 2 (Fig. 4F). Also, likely a result of their higher instantaneous speeds. Collectively, these results suggest that for blebbing cells within microchannels, focal adhesions may functionally inhibit migration.

**Figure 4.**
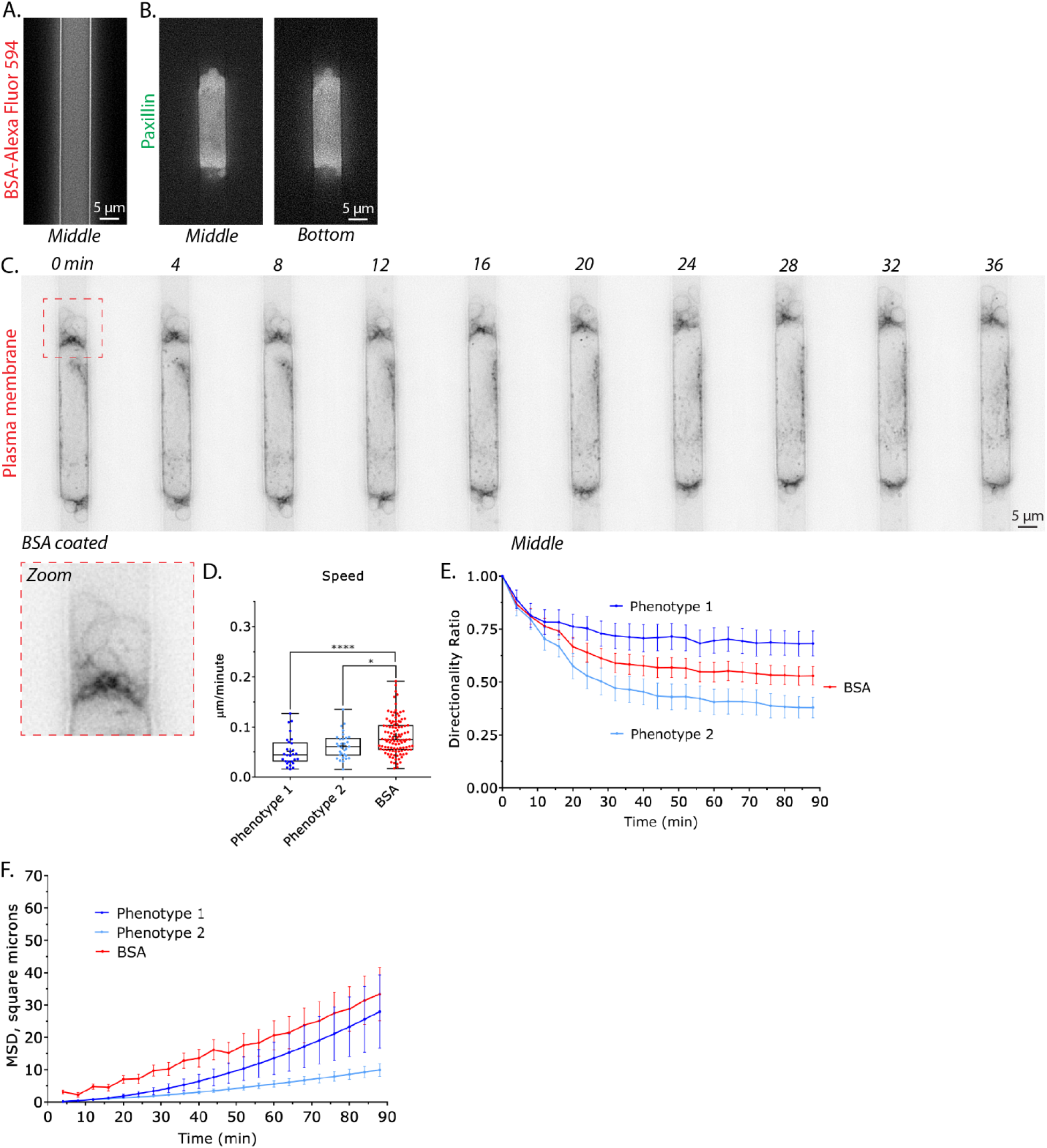
Blebbing cells in poorly adherent microchannels display increased migration parameters. **A.** Confirmation that microchannels are coated with BSA, as indicated by fluorescence from Alexa Fluor 594 conjugated BSA (BSA-Alexa Fluor 594). **B.** Confirmation that a melanoma A375-M2 cell with the focal adhesion marker, EGFP-paxillin, does not form focal adhesions within a BSA coated (1%) microchannel. Middle (*left*) and bottom (*right*) focal planes are shown. **C.** Montage of a A375-M2 cell stained with a far red membrane dye, within a BSA coated microchannel. Zoom shows prominent blebs at the cell leading edge. **D.** A comparison of instantaneous speeds for phenotype 1, 2, and for cells within BSA coated microchannels. Statistical significance was determined by a multiple-comparison test post-hoc. **E.** A comparison of directionality ratio over time for phenotype 1, 2, and for cells within BSA coated microchannels. Error is SEM. **F.** A comparison of Mean Square Displacement (MSD; square microns) over time for phenotype 1, 2, and for cells within BSA coated microchannels. All data are representative of at least three independent experiments. * - p ≤ 0.05, ** - p ≤ 0.01, *** - p ≤ 0.001, and **** - p ≤ 0.0001

### Pharmacologically inhibiting Arp2/3 promotes a fast filopodial-based mode of migration

In previous work using 2D confinement approaches, inhibiting Arp2/3 activity with CK-666 (100 μM) promotes Leader Bleb-Based Migration (LBBM)^6^. The mechanism of which is thought to involve the inhibition of lamellipodia, which requires Arp2/3, and a transition from a poorly to highly contractile (linear formin generated) actin network^16^. Therefore, we wondered if CK-666 treatment would have a similar effect on cells within fibronectin coated (10 μg/mL) microchannels. To our surprise, cells treated with CK-666 (100 μM) no longer formed blebs but instead displayed numerous filopodia at their leading edge (Fig. 5A; *zoom* & Movie S5). Moreover, these cells were fast, having a nearly 2-fold increase in instantaneous speeds (relative to phenotype 1 & 2; Fig. 5B). Directionality over time was between that of phenotype 1 & 2, whereas MSD over time was significantly greater than either phenotype (Fig. 5C-D). Altogether, our results highlight the remarkable diversity of mechanisms melanoma cells have at their disposal for migration within 2D and 3D confined environments.

**Figure 5.**
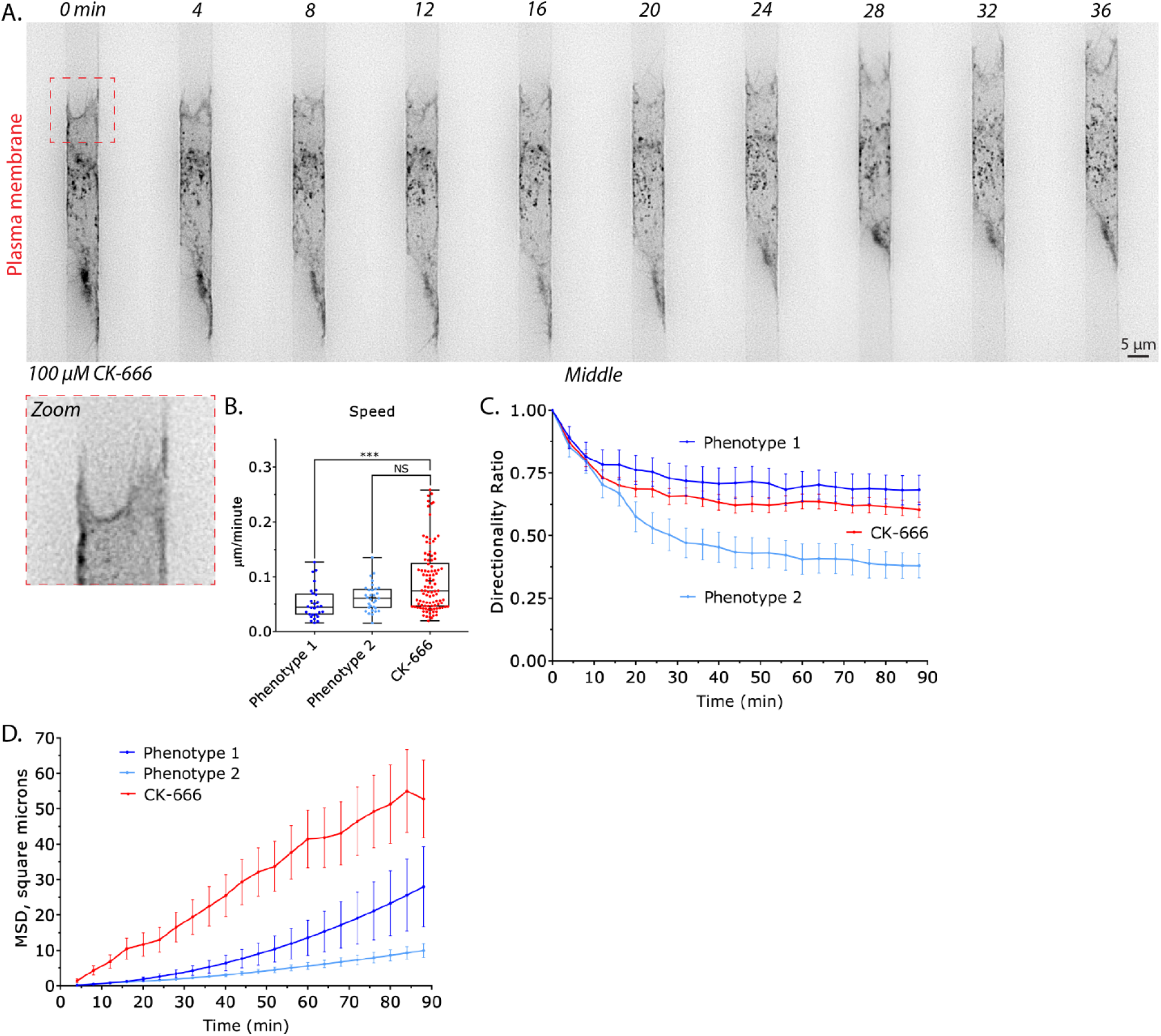
Upon down-regulating Arp2/3 activity, melanoma cells adopt a fast filopodial-based mode of migration. Montage of a melanoma A375-M2 cell treated with the Arp2/3 inhibitor, CK-666 (100 μM), within a fibronectin coated (10 μg/mL) microchannel. Cells were visualized using a far red membrane dye. Zoom shows several filopodia at the cell leading edge. **B.** A comparison of instantaneous speeds for phenotype 1, 2, and CK-666 treated cells. Statistical significance was determined by a multiple-comparison test post-hoc. **C.** A comparison of directionality ratio over time for phenotype 1, 2, and CK-666 treated cells. Error is SEM. **D.** A comparison of Mean Square Displacement (MSD; square microns) over time for phenotype 1, 2, and CK-666 treated cells. All data are representative of at least three independent experiments. * - p ≤ 0.05, ** - p ≤ 0.01, *** - p ≤ 0.001, and **** - p ≤ 0.0001

## Discussion

Several recent studies have shown that cells can utilize a number of different mechanisms for navigating tissues^1^. For melanoma and other cancers, this plasticity is likely to be required to progress from a local to systemic disease. Therefore, through targeting cell migration plasticity, cancer metastasis may be abated or prevented. Under 2D confinement, melanoma and several other cancer cell types have been shown to undergo a phenotypic transition to LBBM. This mode of motility is characterized by the formation of a large and stable bleb at the cell front^5^. In contrast, under conditions of 3D confinement within microchannels, we observe melanoma cells adopting features of both mesenchymal and amoeboid migration.

Within fibronectin coated channels, we found that melanoma cells both adhere and form numerous dynamic blebs. Using the Src family kinase inhibitor, Dasatinib, we could selectively inhibit the formation of focal adhesions^6^. Using this approach, blebbing cells displayed increased migration parameters. Similarly, when placed in microchannels passivated with BSA, cells could not form focal adhesions and migrated with increased instantaneous speeds and directionality over phenotype 1 & 2. Thus, we speculate that focal adhesions restrain the movement of blebbing cells.

Collectively, our results highlight the diversity of mechanisms melanoma cells have at their disposal for migration. For instance, using the Arp2/3 inhibitor, CK-666, we found that cells pre-dominantly adopt a fast filopodial-based mode of migration. This result is also in contrast to what we and others have previously observed under 2D confinement, where treatment with CK-666 promotes LBBM^6,16^. Thus, in contrast to remodeling the actin network in a way favorable to LBBM, down-regulating Arp2/3 promotes the formation of filopodia when melanoma cells are confined within microchannels.

However, these results come with several caveats, which includes the fact that we have only tested one set of channel dimensions. A height and width of 8 μm was chosen on the basis that below 7 μm, the deformation of the nucleus is likely to become a rate limiting factor to migration^14,15^. However, two recent reports showed that cell confinement is sensed by the nuclear envelope, promoting migration through confined environments^17,18^. Therefore, in narrower channels melanoma cells may in fact display a unique set of motile behaviors. Moreover, the topography of the microchannel walls was recently shown to regulate the migration of non-adherent immune cells^19^. A benefit to using microchannels is that they may be used to re-capitulate aspects of the tumor micro-environment, such as an innervating micro-vasculature. Indeed, the lumen of micro-vessels/lymphatics ranges between 1 – 10 μm. Although here we have simulated channel-like environments in general, it remains possible that the introduction of channel coatings, such as VCAM-1 (which decorates micro-vessel walls), may also regulate the motile behavior of melanoma cells. Additionally, it has been shown that the tumor micro-vasculature is under a “pathological state of coagulation.” In fact, the introduction of thrombin, which activates PAR-1 signaling, was recently shown to promote blebbing in breast cancer cells^20^. Thus, in terms of how they may regulate the motile behavior of melanoma cells, several aspects of 3D confinement within microchannels remain to be addressed.

In conclusion, melanoma and other cancer cell types are likely to take advantage of a range of mechanisms for navigating tissues. Thus, the successful abatement or prevention of metastasis necessitates addressing a range of migration modalities.

## Supplemental Information

Supplemental information includes 5 movies and can be found with this article online.

## Methods

### Microchannel design

Our design was based on Bergert and colleagues, having groups of 100 microchannels (100 μm long, 8 μm height, and 8 μm width) to optimize time-lapse imaging^4^. Microchannels were fabricated using standard photo− and soft-lithography procedures.

### Microchannel preparation

PDMS (cat no. 24236-10; Electron Microscopy Sciences, Hatfield, PA) was prepared using a 1:7 ratio of base and curing agent. Uncured PDMS was poured over the wafer mold, placed in a vacuum chamber to remove bubbles, moved to a 37 °C incubator, and left to cure overnight. After curing, small PDMS slabs with microchannels were cut using a scalpel, whereas cell loading ports were cut using a 0.4 cm hole punch (cat no. 12-460-409; Fisher Scientific, Hampton, NH).

For making PDMS coated cover glass (cat no. 12-545-81; Fisher Scientific), 30 μL of uncured PDMS was pipetted at the center of the cover glass, placed in a modified mini-centrifuge, and spun for 30 sec for even spreading. The PDMS coated cover glass was then cured for at least 1 hr on a 95 °C hot plate.

Prior to slab and coated cover glass joining, PDMS surfaces were activated by plasma treatment (cat no. PDC-001-HP; Harrick Plasma, Ithaca, NY). Immediately after activation, slabs were bonded to coated cover glass. For complete bonding, the apparatus was incubated at 37 °C for at least 1 hr.

### Microchannel coating

Prior to microchannel coating, surfaces were first activated by plasma treatment. Fibronectin (cat no. PHE0023; Thermo Fisher) or BSA (cat no. VWRV0332; VWR, Radnor, PA) was used at 10 μg/mL and 1%, respectively, in PBS. Immediately after plasma treatment, fibronectin or BSA solution was pumped into microchannels using a modified motorized pipette. To remove any bubbles pumped into microchannels, the apparatus was left to coat in a vacuum chamber for at least 1 hr. Afterward, fibronectin or BSA solution was aspirated out and microchannels were rinsed twice by pumping in PBS. Finally, microchannels were incubated in complete media over-night at 4 °C before use.

### Microchannel loading

Prior to cells being loaded into microchannels, complete media was aspirated, and microchannels were placed into an interchangeable cover-glass dish (cat no. 190310-35; Bioptechs, Butler, PA). Freshly trypsinized cells in 300 μL of complete media, stained with 1 μL far red membrane dye (cat no. C10046; Thermo Fisher), were pumped into microchannels using a modified motorized pipette. Once at least 20 cells are observed in microchannels by low magnification brightfield imaging, microchannels were covered with 2 mL of complete media. Before imaging, a lid was placed on top the apparatus to prevent evaporation.

### Cell culture

A375-M2 (CRL-3223) were obtained from the American Type Culture Collection (ATCC; Manassas, VA). Cells were cultured in high-glucose DMEM supplemented with 10% FBS (cat no. 12106C; Sigma Aldrich, St. Louis, MO), GlutaMAX (Thermo Fisher), antibiotic-antimycotic (Thermo Fisher), and 20 mM HEPES at pH 7.4 for up to 30 passages.

### Plasmids and transfection

EGFP-paxillin (no. 15233; a gift from Dr. Rick Horwitz) and EGFP-RLC (no. 35680; a gift from Dr. Tom Egelhoff) were obtained from Addgene (Watertown, MA). 1-2 μg of plasmid was used to transfect 500,000 cells in each well of a 6-well plate using Lipofectamine 2000 (5-10 μL; Thermo Fisher) in OptiMEM (400 μL; Thermo Fisher). After 20 min at room temperature, plasmid in Lipofectamine 2000/OptiMEM was then incubated with cells in complete media (2 mL) overnight.

### Pharmacological treatments

Dasatinib (cat no. 6793) and CK-666 (cat no. 3950) were purchased from Tocris Bioscience (Bristol, UK). DMSO (Sigma Aldrich) was used to make 5 mM and 100 μM stock solutions of Dasatinib and CK-666, respectively. Prior to microchannel loading, cells were treated with drug for 1 hr. Simultaneously, microchannels devices were incubated with drug in complete media for at least 1.5 hr before loading cells.

### Cell migration

To perform cell speed and plot of origin analyses, we used an Excel (Microsoft, Redmond, WA) plugin, DiPer, developed by Gorelik and colleagues and the Fiji plugin, MTrackJ, developed by Erik Meijering for manual tracking^21,22^. For minimizing positional error, cells were tracked every other frame. Brightfield imaging was used to confirm that debris were not obstructing the path of a cell.

### Microscopy

Live high-resolution imaging was performed using a General Electric (Boston, MA) DeltaVision Elite imaging system mounted on an Olympus (Japan) IX71 stand with a computerized stage, environment chamber (heat, CO_2_, and humidifier), ultrafast solid-state illumination with excitation/emission filter sets for DAPI, CFP, GFP, YFP, and Cy5, critical illumination, Olympus PlanApo N 60X/1.42 NA DIC (oil) objective, Photometrics (Tucson, AZ) CoolSNAP HQ2 camera, proprietary constrained iterative deconvolution, and vibration isolation table.

### Statistics

All box plots are Tukey in which “+” and line denote the mean and median, respectively. Sample sizes were determined empirically and based on saturation. As noted in each figure legend, statistical significance was determined by either a two-tailed Student’s t-test or multiple-comparison test post-hoc. Normality was determined by a D’Agostino & Pearson test in GraphPad Prism. * - p ≤ 0.05, ** - p ≤ 0.01, *** - p ≤ 0.001, and **** - p ≤ 0.0001

### Data availability

The data that support the findings of this study are available from the corresponding author, J.S.L., upon reasonable request.

## Acknowledgements

We thank members of the Logue Lab for insightful discussions and especially, Yanxiang Deng (RPI), for his help with microchannel design and fabrication. This work was supported by start-up funds from the Albany Medical College, a Young Investigator Award from the Melanoma Research Alliance (MRA; award no. 688232), and a Cancer Research Scholar Grant from the American Cancer Society (ACS; award no. RSG-20-019-01 - CCG) to J.S.L.

## Author Contributions

J.S.L. conceived and designed the study. S.R.G. performed all laboratory research. A.J.C. assisted with photo− and soft-lithography procedures. J.S.L. wrote the manuscript with comments from S.R.G.

## Competing Financial Interests

The authors declare no competing financial interests.

## Notes

### Competing Interest Statement

The authors have declared no competing interest.

