## Supplemental Information for "Melanoma cells adopt features of both mesenchymal and amoeboid migration within confining channels"

^4^Present address: School of Biomedical Engineering, Korea University, 02841, Seoul, Republic of Korea

^5^Present address: Interdisciplinary Program in Precision Public Health, Korea University, 02841 Seoul, Republic of Korea

*Corresponding author

Short title: melanoma cell migration in confining channels

**Supplemental Movies**

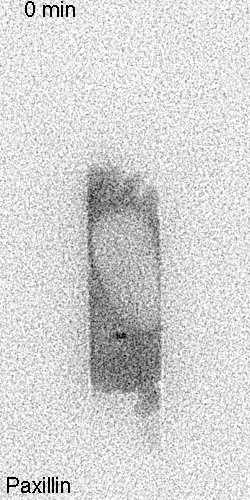

**Supplemental movie 1.** Time-lapse imaging of a melanoma A375-M2 cell (*phenotype 1*) within a fibronectin coated (10 µg/mL) microchannel with the focal adhesion marker, EGFP-paxillin.

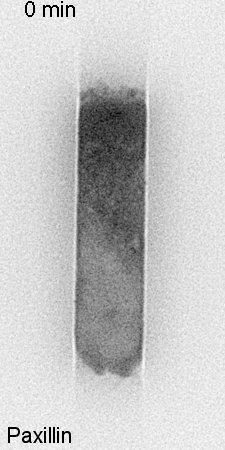

**Supplemental movie 2.** Time-lapse imaging of a melanoma A375-M2 cell (*phenotype 2*) within a fibronectin coated (10 µg/mL) microchannel with the focal adhesion marker, EGFP-paxillin.

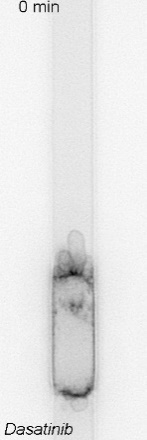

**Supplemental movie 3.** Time-lapse imaging of a melanoma A375-M2 cell within a fibronectin coated (10 µg/mL) microchannel treated with the Src family kinase inhibitor, Dasatinib (10 µM). Cells were stained with a far red membrane dye.

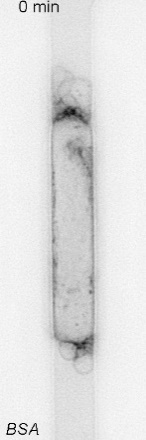

**Supplemental movie 4.** Time-lapse imaging of a melanoma A375-M2 cell within a BSA coated (1%) microchannel. Cells were stained with a far red membrane dye.

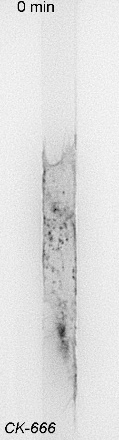

**Supplemental movie 5.** Time-lapse imaging of a melanoma A375-M2 cell within a fibronectin coated (10 µg/mL) microchannel treated with the Arp2/3 inhibitor, CK-666 (100 µM). Cells were stained with a far red membrane dye.
